# Opposing effects of prostaglandin receptor EP2 signaling in mouse immune cells and neurons after status epilepticus

**DOI:** 10.1101/395566

**Authors:** Nicholas H. Varvel, Di Chen, Ariel Biegel, Raymond Dingledine

## Abstract

A multidimensional inflammatory response ensues after status epilepticus (SE), driven partly by cyclooxygenase-2 mediated activation of prostaglandin EP2 receptors. Here we identify those effects of EP2 antagonism that are reproduced by conditional ablation of EP2 receptors in immune myeloid cells and forebrain neurons. Removal of myeloid cell EP2 dampened induction of hippocampal IL-6, prevented erosion of the blood-brain barrier, accelerated weight regain, and relieved ptosis following SE. Elimination of EP2 receptor from neurons enhanced neuronal injury, elevated hippocampal induction of the pro-inflammatory cytokines, TNFα and Ccl2, promoted deterioration of the blood-brain barrier, delayed weight recovery, and worsened animal posture and activity. Taken together these data highlight the complexities in neuroinflammatory signaling, wherein activation of EP2 receptors in innate immune cells is deleterious but neuronal EP2 signaling is protective. Effective treatments targeting brain prostaglandin signaling pathways should be cell targeted to be optimally effective.

**Significance Statement:** Seizures reduce quality of life, promote the development of epilepsy, and can be fatal. We previously identified inflammation, via prostaglandin receptor EP2 activation, as a driver of undesirable seizure-induced effects. However, the relevant EP2-expressing cell types remain unclear. We identify innate immune cells as a driver of the EP2-related negative consequences of seizures, whereas neuronal EP2 signaling is protective. Genetic removal of EP2 from immune cells was beneficial, accelerating weight gain and limiting behavioral deficits. Elimination of EP2 from neurons was harmful, worsening behavioral deficits and promoting neuronal damage. These findings enhance our understanding of the complex inflammatory processes engaged after seizures and will assist in the development of beneficial therapies to enhance quality of life in individuals susceptible to seizures.

## Introduction

In both man and rodents, the extended seizures of status epilepticus (SE) can result in marked brain inflammation, temporary opening of the blood-brain barrier (BBB), and neuronal damage in select brain regions leading to substantial decline in quality of life. Patients in refractory SE typically show elevated levels of chemokines and cytokines, such as IL-6, IL-8, TNFα, and CXCL10 in cerebrospinal fluid (1). Examination of the temporal cortex in a patient with new-onset focal seizures that progressed into refractory SE revealed robust activation of astrocytes and microglia (2).

In rodent hippocampus, several inflammatory mediators – cyclooxygenase-2 (Cox-2), Ccl2, and IL-1β – are strongly induced within 30 minutes of SE onset, and the induction of other cytokines follows (3). Interestingly, inhibiting inflammation after SE can prevent transient erosion of the BBB, provide neuronal protection, alleviate behavioral morbidity, and rescue subsequent behavioral deficits in rodents (4-7). Our previous studies have identified activation of the prostanoid receptor EP2 as a strong driver of neuroinflammation, mortality, and behavioral deficits following SE. Administration of a brain-permeable EP2 antagonist starting 2-4 hours after SE onset produces a broad range of beneficial effects including reduction in delayed mortality, blunted neuroinflammation, prevention of BBB opening, neuroprotection, accelerated weight regain, and cognitive improvement (3, 8-10). However, in these studies EP2 receptors were globally inhibited; thus, the relevant cell types involved in EP2-driven consequences of SE remain enigmatic.

In the present studies we generated two cell type-specific EP2 conditional knockout lines. In the one mouse line, we genetically ablated the EP2 gene from CD11b-expressing innate immune myeloid cells (microglia and blood monocytes), and we compared the SE-induced phenotypes of myeloid EP2 conditional knockout mice (McKO) to their EP2-sufficient gender-matched littermate controls (mEP2+) to determine which beneficial effects of EP2 antagonism can be attributed to EP2 activation in innate immune myeloid cells. In the second mouse line, EP2 was genetically ablated from synapsin 1-expressing forebrain neurons, and we compared the SE-induced phenotypes of neuronal EP2 conditional knockout mice (NcKO) to their EP2-sufficient gender-matched littermate controls (nEP2+) to determine which beneficial effects of EP2 antagonism can be attributed to EP2 activation in forebrain neurons.

The results point to a deleterious action of myeloid cell signaling and a protective effect of neuronal EP2 activation after SE, highlighting, for the first time, a cell-type specific dichotomous role of EP2 signaling after SE.

## Results

### McKO mice display less behavioral deterioration following status epilepticus

Previously we have shown that during the first 24-48 hours following SE, rodents lose 10-20% of their body weight and then slowly begin to recover in the following days (8, 9, 11). In the current study both mEP2+ and McKO mice lost ∼9% of their body weight in the 24 hours after SE onset. By the second day mEP2+ mice continued to lose weight, and they didn’t recover their lost body weight, such that by the fourth day the mEP2+ animals on average had lost ∼15% of their original body weight (Figure 1A). In contrast, McKO mice began to regain their lost weight by the third day. By the fourth day the McKO had regained all but ∼5% of their original body weight prior to SE. We compared the mean area under the curve (AUC) between days 1 and 4 for statistical analysis. The mean AUC was significantly larger for mEP2+ mice (42.8± 5.36 % weight loss x day, mean± SEM) than for McKO mice (27.4± 3.50 %) (Figure 1A, p=0.03, unpaired *t* test).

**Figure 1.**
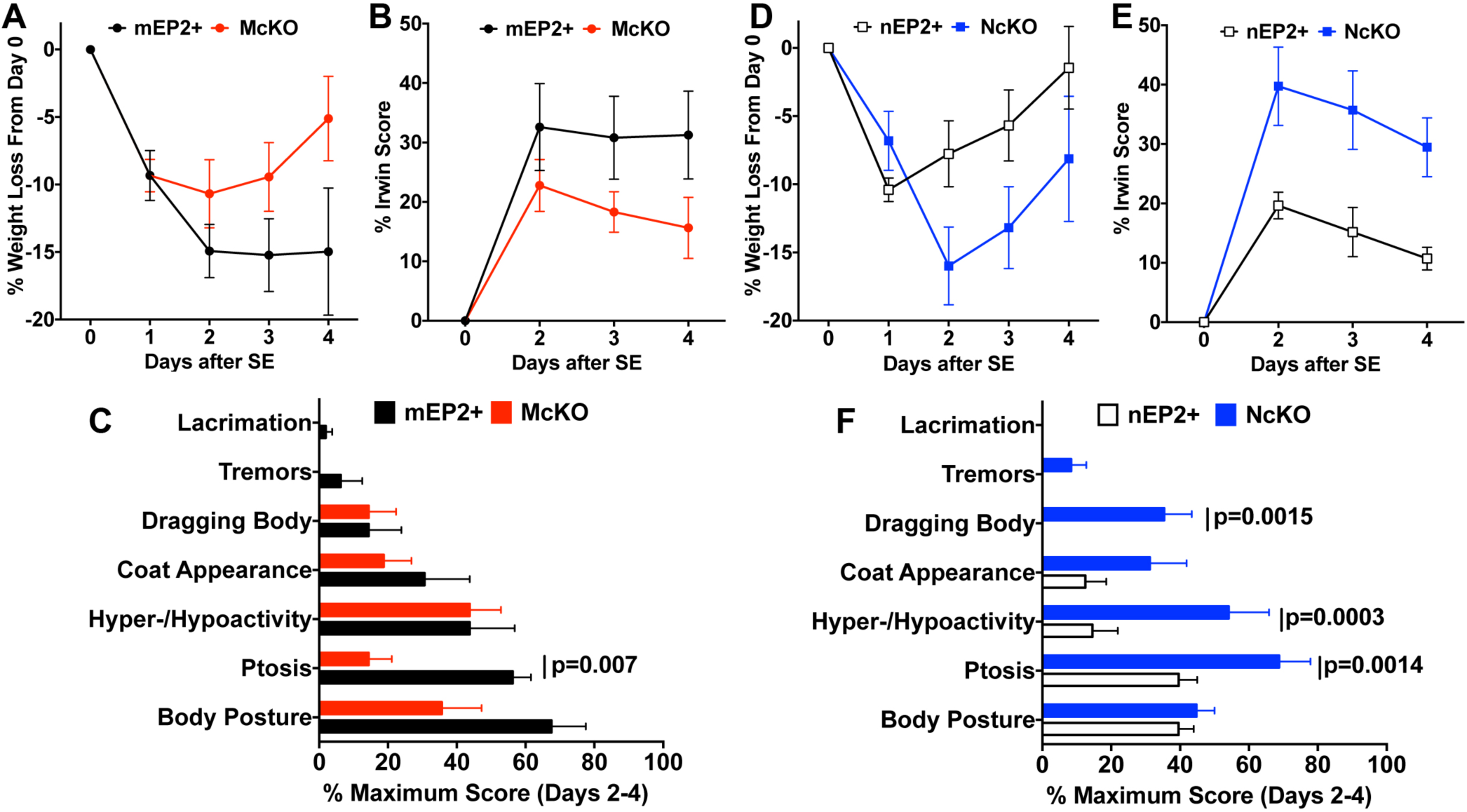
Accelerated weight gain and improved Irwin scores in McKO mice, but not in NcKO mice, after SE. (A) Effect of EP2 deficiency in myeloid cells on mouse body weight change after SE (n=8/day in both mEP2+ (black circles) and EP2 McKO (red circles). AUC was smaller for the McKO group on days 1-4, p=0.03, unpaired t-test. (B) Comparison of the modified Irwin scores taken on days 2-4 did not reveal a difference between mEP2+ (open circles) and McKO (red circles), AUC p=0.12, unpaired t-test. Data are shown as a percent of the maximum possible score, with 0 indicating a normal, healthy mouse. (C) Subdividing the mean Irwin scores for days 2-4 into each metric revealed a significant difference in ptosis scores between mEP2+ (n=8, black bars) and McKO (n=8, red bars) mice, but no significant differences in the other behavioral measures were encountered by ANOVA with post-hoc Sidak for selected pairs. Data are shown as a percent of the maximum score. (D) Effect of EP2 deficiency in neurons on mouse body weight change (n=8/day in both nEP2+ (open squares) and EP2 NcKO (blue squares). AUC was not different between nEP2+ and NcKO mice on days 1-4, p=0.23, unpaired t-test. (E) Comparison of the modified Irwin scores taken on days 2-4 revealed worse scores in nEP2+ (open squares) compared to NcKO (blue squares), AUC p=0.01, unpaired t-test. Data are shown as a percent of the maximum possible score. (F) Subdividing the mean Irwin scores for days 2-4 into each metric revealed a significant difference in ptosis, hyper-/hypoactivity, and dragging body scores between nEP2+ (n=8, open bars) and NcKO (n=8, blue bars) mice, but no significant differences in the other behavioral measures were encountered by ANOVA with post-hoc Sidak for selected pairs. Data are shown as a percent of the maximum score. Bars represent mean percent + SEM.

Using a modified Irwin test we determined the effect of eliminating EP2 from innate immune myeloid cells on a selected subset of normal behavioral features. The data were plotted as a percent of the maximum possible score for each genotype following SE; higher scores reflect more functional impairment. In general, modified Irwin scores were high 24 hours after SE and then declined towards normal regardless of genotype. mEP2+ mice showed higher modified Irwin scores than McKO mice, suggesting more deterioration in normal physiological characteristics in EP2-sufficient mice. The mean± SEM AUC for mEP2+ mice was 62.7± 13.7 % Irwin score x day, compared to 37.5± 8.84 % for McKO mice (Figure 1B, p=0.12 by unpaired *t* test). We also found that relief from ptosis had a dominant contribution to the observed behavioral differences between genotypes (Figure 1C). Taken together, these findings indicate that McKO mice show less behavioral deficit, with ptosis dominating this observation, compared to their littermate mEP2+ counterparts in the days following SE, suggesting that myeloid cell EP2 signaling has a deleterious role in mouse well-being after SE.

### The two control EP2+ strains display different responses to pilocarpine

The temporal change in percent weight loss after pilocarpine-induced SE in mEP2+ (Figure 1A) was similar to the percent weight loss we have previously encountered in stock C57BL/6 mice obtained from Charles River (3, 8). However, forthcoming results will show that nEP2+ mice do not respond in similar fashion to pilocarpine as stock C57BL/6 mice and mEP2+ mice. About one third of the experiments utilizing the two cKO lines were performed concurrently, and the mice were housed in the same room so environmental differences are unlikely to fully account for the dissimilar responses to pilocarpine. However, we cannot rule out genetic differences in the two strains as a result of spontaneous mutations or inadvertent introduction of genetic material during extensive backcrossing to C57BL/6 stock mice. Nonetheless, each EP2+ line (myeloid and neuronal) is the appropriate comparison strain for their conditional knockout counterparts, and this comparison of the two EP2-sufficient lines demonstrates the value of using littermate controls for genetically manipulated mice.

### NcKO mice display enhanced behavioral deterioration following status epilepticus

In the 24 hours after SE, NcKO mice lost ∼7% of their body weight, whereas their nEP2+ littermates lost nearly 11% of their initial body weight. By the second day the nEP2+ mice had begun to regain their lost weight such that by the fourth day after SE the nEP2+ controls had nearly regained their SE-induced lost weight (Figure 1D). In contrast, NcKO mice continued to lose weight through the second day after SE such that the NcKO mice had lost nearly 16% of their body weight 48 hours after SE. By the fourth day, the NcKO mice were still down over 8% of their initial body weight (Figure 1D). The mean AUC between days 2 and 4 was not statistically different for nEP2+ (25.5± 4.76 % weight loss x day, mean± SEM) and NcKO mice (35.7± 6.54 %) (Figure 1D, p=0.23)

Eliminating EP2 from neurons resulted in higher Irwin scores when compared to nEP2+ littermates, suggesting more deterioration in normal physiological characteristics in the absence of neuronal EP2. The mean± SEM AUC for nEP2+ mice was 30.3± 5.7 % Irwin score x day, compared with 69.9± 12.0 % for NcKO mice (Figure 1E, p=0.01 by unpaired *t* test). We also found worsened ptosis, abnormal activity, and dragging of body in NcKO mice compared to nEP2+ controls (Figure 1F). Taken together, these findings indicate that NcKO mice show enhanced behavioral deficit, with body dragging, general activity, and ptosis dominating this observation, suggesting that neuronal EP2 signaling has a beneficial role in mouse behavior after SE, in contrast to myeloid cell EP2 signaling.

### Serum albumin levels in the cortex are elevated after status epilepticus in mEP2+ mice, but not in McKO mice

Insults to the brain, including SE, can degrade the BBB, leading to extravasation of blood-derived proteins such as serum albumin. The ensuing penetration of albumin into brain parenchyma has been purported to trigger epileptogenesis (12, 13). Serum albumin was detected in cortical homogenates obtained from mEP2+ mice perfused with PBS four days following pilocarpine-induced SE, compared with minimal albumin detected in cortex of saline-treated mEP2+ control mice (Figure 2A). In contrast, about half of the McKO mice had elevated albumin in the cortex in both saline and pilocarpine groups, and there was no further increase in cortices from pilocarpine-treated McKO mice compared to saline-treated McKO controls (Figure 2). These findings suggest that myeloid EP2 receptors influence the integrity of the BBB under normal conditions, and that EP2 signaling in innate immune myeloid cells enhances erosion of the BBB after pilocarpine-induced SE.

**Figure 2.**
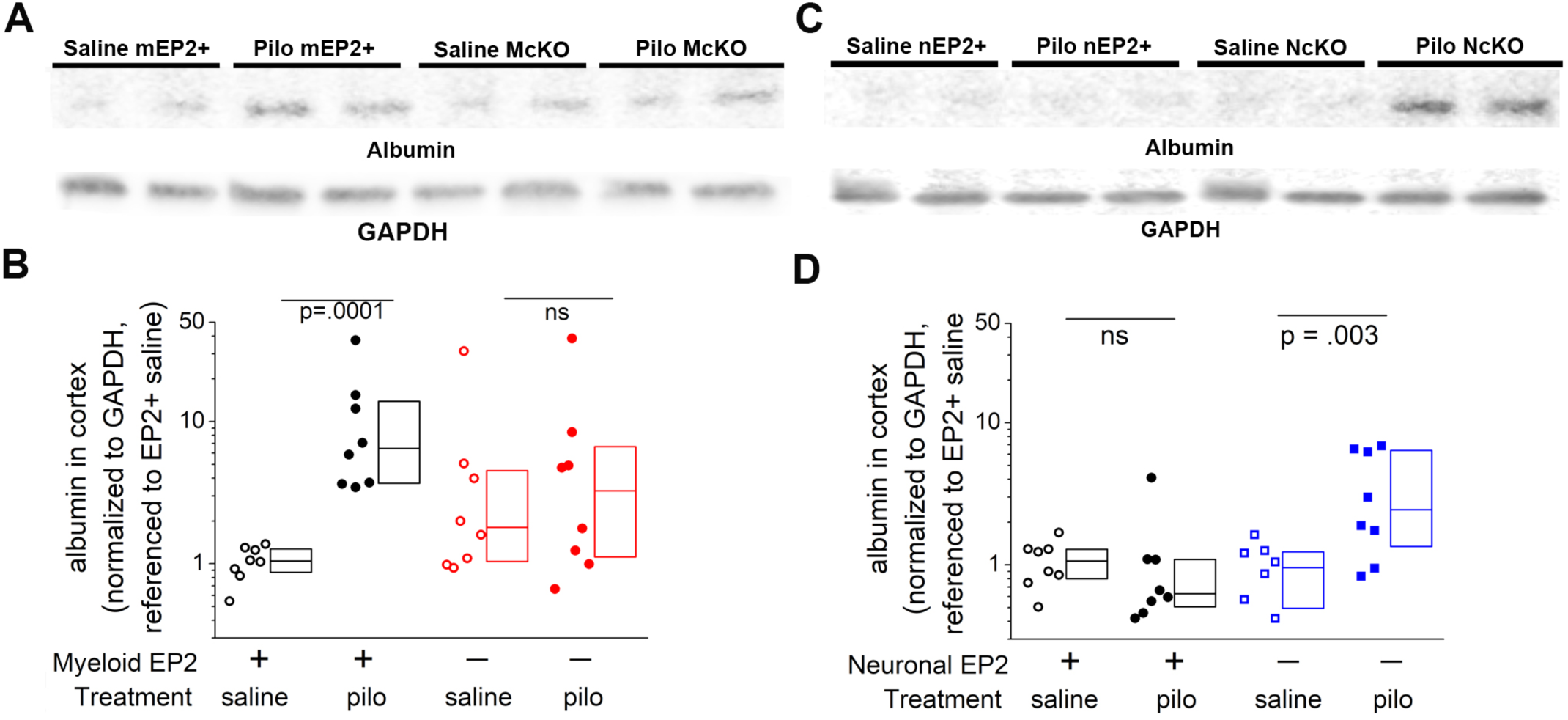
Serum albumin extravasation into the brain 4 days after SE was used to evaluate the integrity of the BBB. (A) Albumin protein levels in cortical homogenates of saline-treated and pilocarpine-treated mEP2+ and McKO mice were measured by Western blot with GAPDH as loading control (n=8/group). Two representative samples near the median of each group are shown. (B) Normalized band intensity of the albumin protein relative to that of GAPDH and referenced to the mean albumin/GAPDH ratio in saline treated mEP2+ mice. The average log ratio of albumin was compared between each experimental group by one-way ANOVA with post-hoc Sidak test with selected pairs. Boxes represent the 25^th^/75^th^ percentiles. The horizontal line in the boxes presents the median value. (C) Albumin protein levels in cortical homogenates of saline-treated and pilocarpine-treated nEP2+ and NcKO mice were measured by Western blot with GAPDH as loading control (n=8/group). Two representative samples near the median of each group are shown. (B) Normalized band intensity of the albumin protein relative to that of GAPDH and referenced to the mean albumin/GAPDH ratio in saline treated nEP2+ mice. The average log ratio of albumin was compared between each experimental group by one-way ANOVA with post-hoc Sidak test with selected pairs.

### Serum albumin levels in the cortex are elevated after status epilepticus in NcKO mice, but not in nEP2+ littermates

Serum albumin levels were elevated in cortical homogenates obtained from NcKO mice perfused with PBS four days following pilocarpine-induced SE, compared with minimal albumin detected in cortex of saline-treated nEP2+ control mice (Figure 2C). In contrast to previous studies (8, 11), pilocarpine-treated nEP2+ mice subject to SE four days prior did not show a significant increase in cortical serum albumin levels (Figure 2D). These observations reveal the unexpected finding that neuronal EP2 signaling maintains the integrity of the BBB in the days following pilocarpine SE.

### Neuronal EP2, but not myeloid cell EP2, is neuroprotective after SE

We assessed the degree of neuronal damage four days after pilocarpine-induced SE by examining Fluoro-Jade B staining in hippocampal brain sections. As expected, Fluoro-Jade B staining was encountered in the CA1 and CA3 hippocampal subfields as well as the hilar region of the dentate gyrus in both mEP2+ and McKO mice after SE. Surprisingly, in the CA1 subfield the number of damaged neurons appeared to be similar regardless of genotype (Figure 3A). This impression was confirmed by counting the number of Fluoro-Jade B+ pyramidal neurons in the CA1 and CA3 subfields as well as the hilar neurons. No significant differences in number of damaged neurons were encountered (Figure 3B).

**Figure 3.**
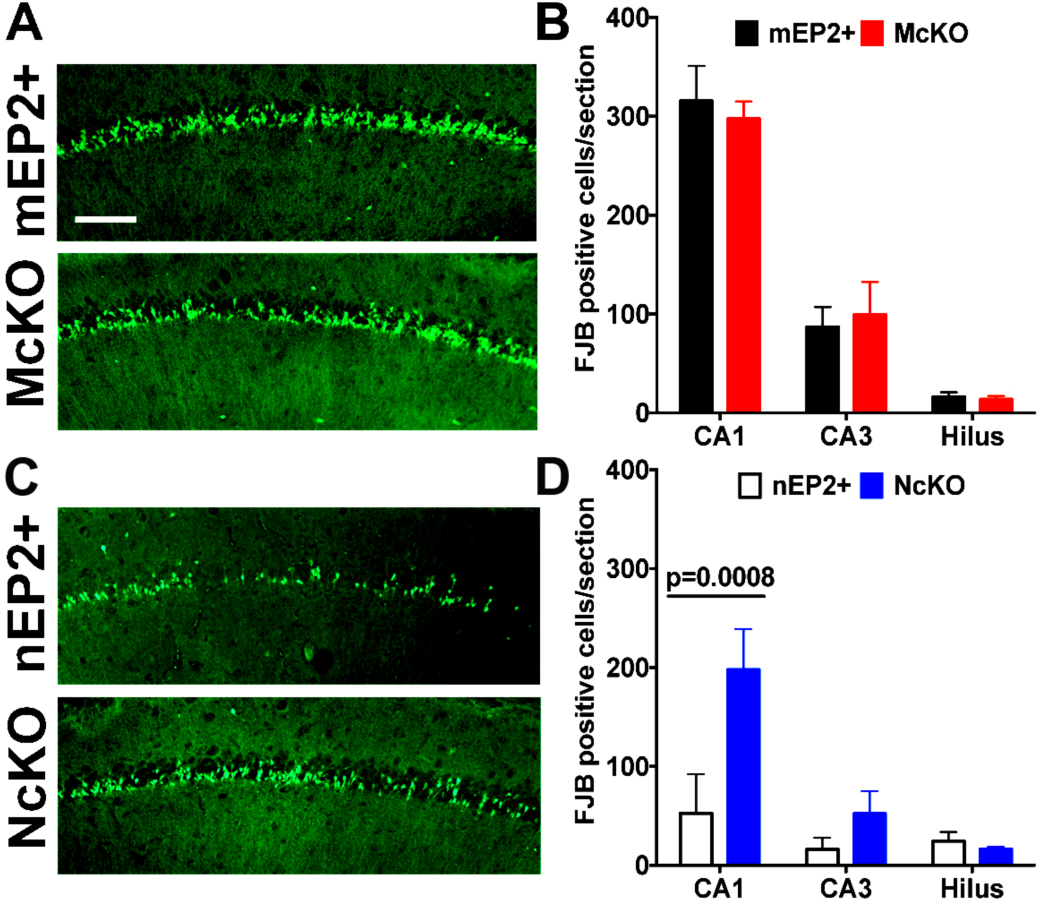
Enhanced neuronal damage is observed in NcKO mice after SE; Neuroprotection is not observed in McKO mice after SE. (A) Fluoro-Jade B staining in hippocampal tissue sections 4 days after SE shows similar numbers of damaged neurons in the CA1 subfield of mEP2+ compared to McKO littermates (Scale bar: 200 μm). (B) a plot of the number of Fluoro-Jade B positive cells in hippocampal cell layers. The bars indicate the mean ± SEM of each group (n=8/group); no significant differences were found by one-way ANOVA with Sidak multiple-comparisons test. (C) Fluoro-Jade B staining in hippocampal tissue sections 4 days after SE shows increased numbers of damaged neurons in the CA1 subfield of NcKO mice compared to nEP2+ littermates (Scale bar: 200 μm). (D) a plot of the number of Fluoro-Jade B+ cells in hippocampal cell layers. The bars indicate the mean ± SEM of each group (n=8/group); significant differences were found by one-way ANOVA with Sidak multiple-comparisons test in CA1 subfield.

Hippocampal neuronal damage was also assessed in NcKO and EP2+ littermates four days after SE. Whereas Fluoro-Jade B staining was observed in both nEP2+ and NcKO tissues (Figure 3C), the number of Fluoro-Jade B reactive neurons was higher in the NcKO mice in the CA1 and CA3 subfields of the hippocampus, reaching statistical significance in the CA1 region (Figure 3D). These findings suggest that neuronal EP2 signaling can be protective after pilocarpine-induced SE.

### Hippocampal induction of IL-6 is selectively blunted in McKO mice after SE

To gain further insight into the role of myeloid cell EP2 signaling after SE, we investigated the expression profile of a panel of inflammatory mediators in hippocampus from EP2+ and McKO mice. Quantitative real-time PCR (qRT-PCR) was performed on hippocampal tissue to measure the levels of inflammatory mediators as well as glial markers previously demonstrated to be induced in the mouse hippocampus after SE (7, 8, 11). The basal mRNA levels of the inflammatory mediators and glial markers were nearly identical in hippocampus obtained from saline treated mEP2+ control mice and McKO animals (Figure 4A). The mRNA levels of the inflammatory mediators IL-1β, TNFα, Ccl2, and Cox-2 were all induced in mEP2+ and McKO mice at similar levels, although variability in TNFα and Ccl2 mRNA levels could have masked a 50-70% suppression. However, the induction of hippocampal IL-6 was almost completely prevented in the McKO mice subject to pilocarpine-induced SE compared with mEP2+ mice (p=0.0025; Figure 4B). No significant differences were encountered in the glial markers examined between mEP2+ and McKO mice (Figure 4C).

**Figure 4.**
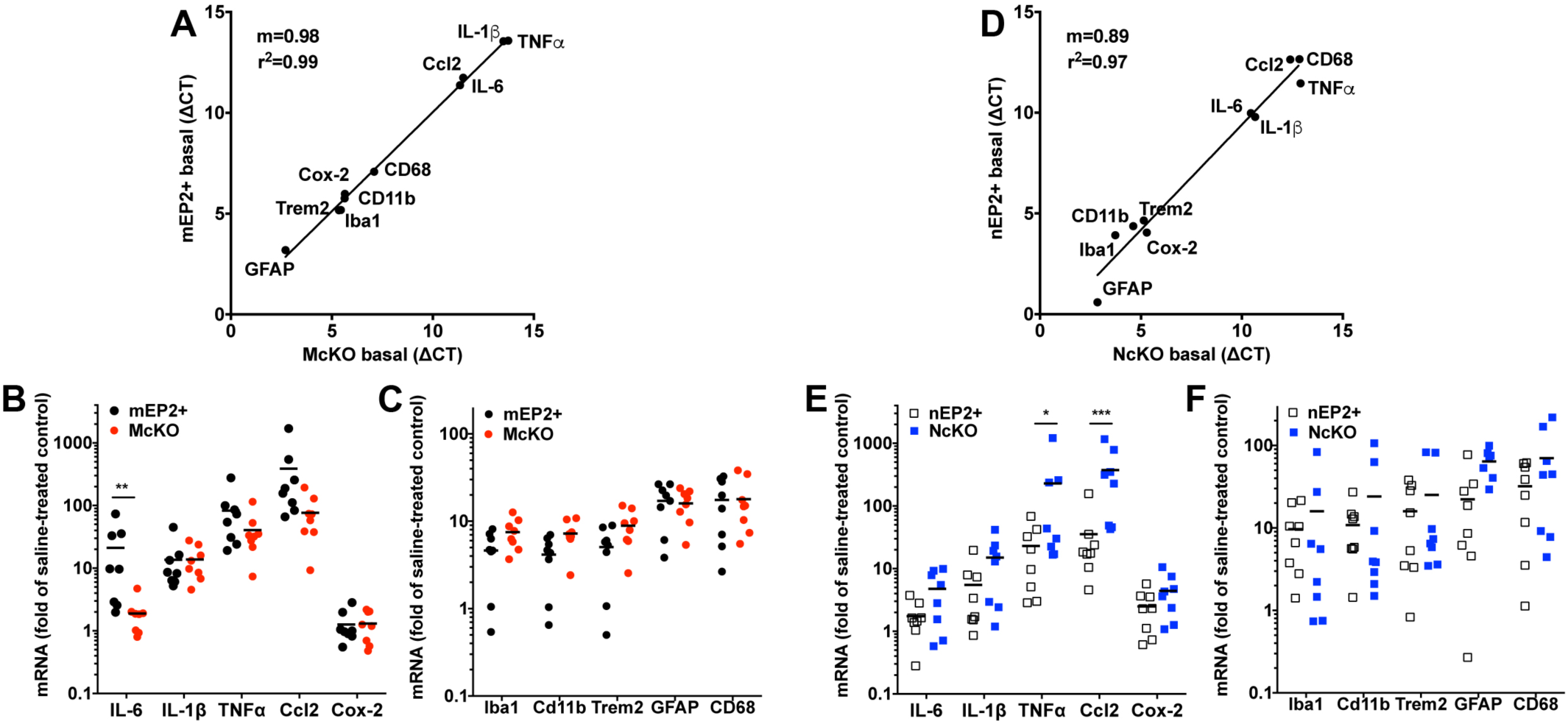
Expression of inflammatory mediators and glial markers in hippocampus 4 days after SE. (A) The basal hippocampal levels of mRNA encoding inflammatory mediators and glial markers were similar in mEP2+ and McKO animals injected with saline solution; each symbol represents the mean ΔCT of 8 mice of both genotypes. Mean fold induction of the inflammatory mediators (B) and glial markers (C) in hippocampus is represented by the horizontal line and is plotted for mEP2+ (n=8) and McKO (n=8) mice 4 days after SE. (D) The basal hippocampal levels of mRNA encoding inflammatory mediators and glial markers were similar in nEP2+ and NcKO animals injected with saline solution; each symbol represents the mean ΔCT of 8 mice of both genotypes. Mean fold induction of the inflammatory mediators (E) and glial markers (F) in hippocampus is represented by the horizontal line and is plotted for nEP2+ (n=8) and NcKO (n=8) mice 4 days after SE. Each symbol represents an individual mouse, with induction referenced to the mean of the appropriate saline solution-treated group (*p=0.0184, **p=0.0025, and ***p=0.0008 by one-way ANOVA with Sidak multiple-comparisons test).

### Hippocampal induction of TNFα and Ccl2 is selectively elevated in NcKO mice after SE

Saline solution-treated nEP2+ and NcKO mice showed nearly identical basal levels of mRNA of the inflammatory mediators and glial markers in the hippocampus (Figure 4D). The mRNA levels of the inflammatory mediators IL-6, IL-1β, and Cox-2 were all induced in nEP2+ control mice and NcKO mice at similar levels, although variability in IL-6 and TNFα could have masked an induction in NcKO hippocampus. TNFα (*p=0.0184, Figure 4E) and CCL2 (***p=0.0008, Figure 4E) mRNA was induced in NcKO hippocampus approximately 10-fold over the induction observed in hippocampus of nEP2+ controls. No significant differences were encountered in the glial markers examined between nEP2+ and NcKO mice, although there were general trends for greater induction in NcKO hippocampus (Figure 4F).

### Systemic Administration of Pilocarpine Induces Typical Status Epilepticus in the Absence of EP2 in Innate Immune Myeloid Cells or Neurons

Upon entering SE, three out of 12 McKO mice died in SE compared with two out of 11 mEP2+ littermate control mice (Figure 5A). Of the 18 surviving mice, one mouse died in each group in the following four days after interruption of SE by diazepam (Figure 5B), indicating that the absence of myeloid cell EP2 did not modify mouse survival during SE or delayed mortality in the days following SE.

**Figure 5.**
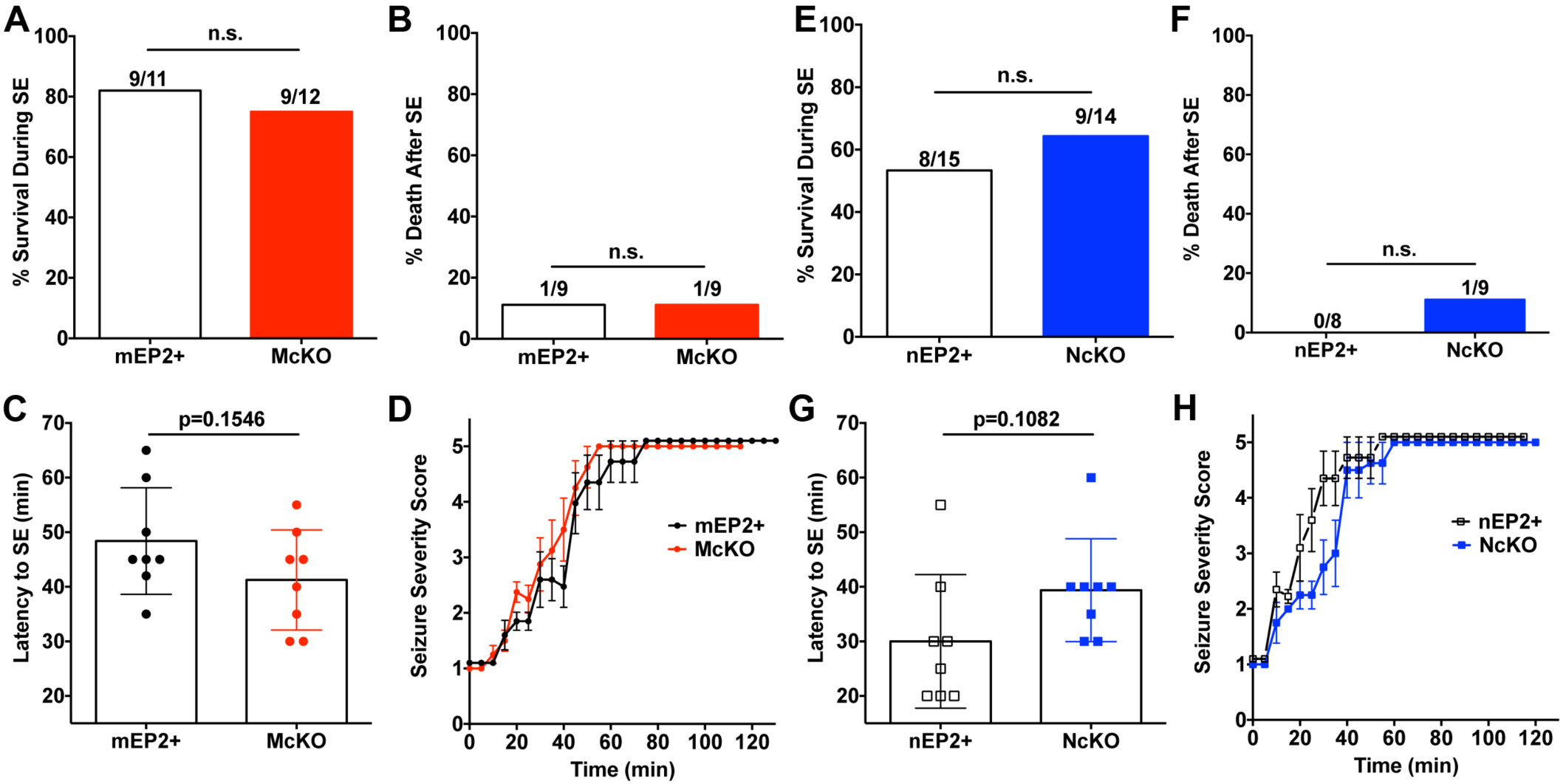
Behavioral seizure intensity and mortality appear similar in McKO mice and littermate controls, and NcKO and littermate controls. (A) The mortality observed during SE was similar in McKO and mEP2+ littermates (Fisher’s exact test). (B) Mortality after SE, defined as deaths that occurred during the 4 days after interruption of SE, was identical between McKO and mEP2+ mice (Fisher’s exact test). (C) The latency to SE as judged by the behavioral seizure score was similar in mEP2+ and McKO mice (n=8/genotype, unpaired *t* test). The bars represent mean ± SEM. (D) The behavioral seizure score was tabulated every 5 min until the seizure was interrupted by diazepam 1 hour after SE onset in mEP2+ and McKO mice (n=8/genotype; EP2+ black circles, McKO red circles). The temporal evolution of SE was similar in both genotypes. The average behavioral scores are represented as mean ± SEM. (E) The mortality observed during SE was similar in NcKO and nEP2+ littermates (Fisher’s exact test). (F) Mortality after SE, defined as deaths that occurred during the 4 days after interruption of SE, was not significantly different between NcKO and nEP2+ mice (Fisher’s exact test). (G) The latency to SE as judged by the behavioral seizure score was similar in nEP2+ and NcKO mice (n=8/genotype, unpaired *t* test). The bars represent mean ± SEM. (H) The behavioral seizure score was tabulated every 5 min until the seizure was interrupted by diazepam 1 hour after SE onset in nEP2+ and NcKO mice (n=8/genotype; nEP2+ open squares, NcKO blue squares). The temporal evolution of SE was similar in both genotypes. The average behavioral scores are represented as mean ± SEM.

Next we evaluated the latency to SE in the 16 mice that survived until the four-day time point. On average the EP2+ sufficient mice entered SE ∼48 minutes after pilocarpine administration, whereas the McKO mice entered SE ∼41 minutes after pilocarpine, a difference not statistically significant (Figure 5C). In addition, the temporal evolution of seizure severity behavior scores was similar between the EP2+ and McKO groups (Figure 5D).

After entering SE five out of 14 NcKO animals died in SE compared with seven out of 15 nEP2+ mice (Figure 5E). In the following four days after SE interruption by diazepam, one of the NcKO mice died (Figure 5F), indicating that the absence of neuronal EP2 did not alter mouse survival during SE or delayed mortality in the days following SE.

Finally we evaluated the latency to SE in the 16 mice that survived until completion of the experiment four days after SE. On average the EP2+ mice entered SE ∼30 minutes after pilocarpine, whereas the NcKO littermates entered SE ∼40 minutes after SE, a difference not statistically significant (Figure 5G). Also, the temporal evolution of seizure severity behavior scores was similar between the nEP2+ and NcKO groups (Figure 5H). Taken together, these findings indicate that the absence of innate immune myeloid cell EP2 or neuronal EP2 does not alter mortality during SE, delayed mortality after SE, or seizure susceptibility. This contrasts sharply with the global EP2 knockout mouse, which is very resistant to entering SE after pilocarpine (not shown).

### Cre recombinase robustly ablates EP2 from CD11b+ immune cells

We did experiments to determine the extent of EP2 ablation in myeloid cells. Because we were unable to validate three different EP2 antibodies with two different EP2 knockout mouse strains (Material and Methods, Figure 7), we sought to determine the mRNA levels of EP2 in CD11b+ splenocytes. Nearly 60,000 CD11b+ cells were obtained from each spleen of three mEP2+ and three McKO mice by flow cytometry sorting (Figure 6A). To assess relative levels of CD11b+ splenocyte EP2 mRNA in both genotypes, the delta cycle threshold value (ΔCt) for EP2 in mEP2+ splenocytes was compared to McKO splenocytes, using CD11b mRNA levels to normalize. The mean ΔCt for EP2+ spleen cells was 0.11, whereas the mean ΔCt for McKO cells was 11.12 (Figure 6B), indicating a near complete loss of EP2 mRNA in McKO CD11b+ splenocytes.

**Figure 6.**
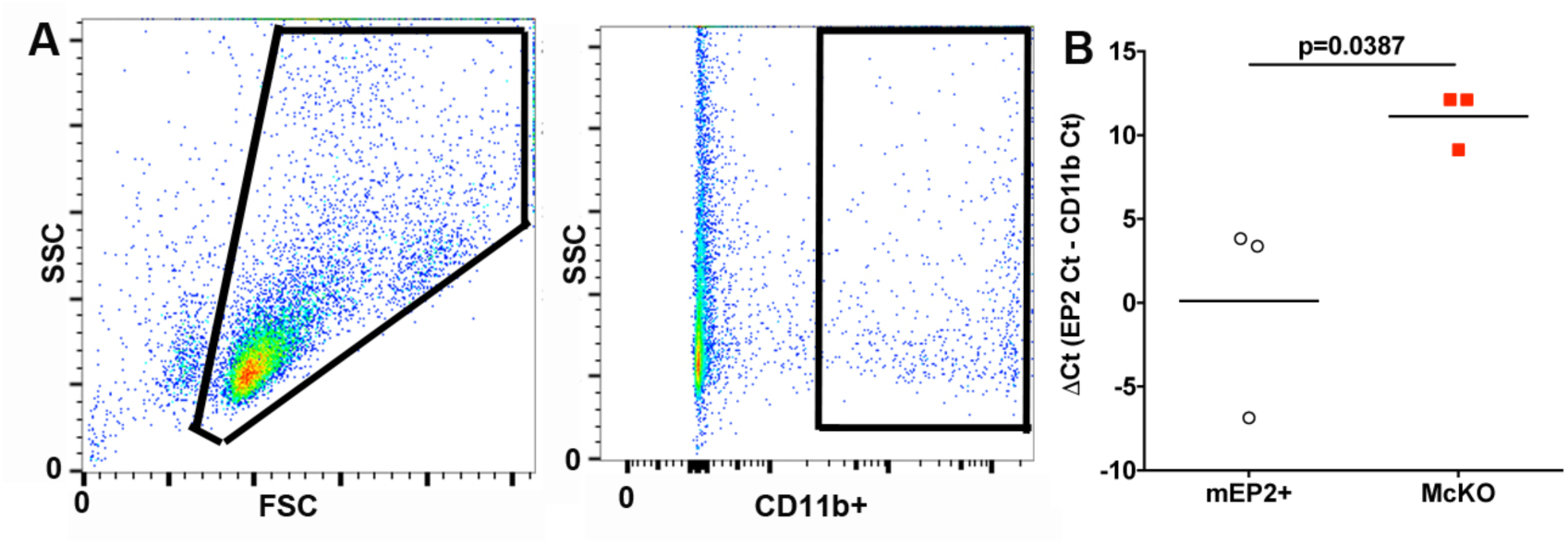
EP2 mRNA is robustly ablated in CD11b+ innate immune cells from McKO mice. (A) Single cell suspensions of CD11b+ spleen cells from three mEP2+ and three McKO mice were gated on CD11b. (B) The mean ΔCt for EP2 is represented by the horizontal line for mEP2+ and McKO mice. Each symbol represents an individual mouse (p=0.0387 by students t-test).

## Discussion

Our studies were designed to identify the effects of systemic pharmacologic EP2 antagonism that are reproduced by conditional ablation of EP2 receptors in either innate immune myeloid cells (microglia and monocytes) or forebrain neurons. We show that ablation of EP2 in microglia and monocytes alleviates many of the deleterious consequences of SE including suppression of weight regain, ptosis, and erosion of the BBB. Interestingly, however, elimination of EP2 in neurons was deleterious, worsening many SE-induced phenotypes, including behavioral deficits, BBB deterioration, and neuronal damage.

A key feature of this study was the use of littermate controls for each of the two EP2 cKO mouse lines. This proved critical during data analysis because comparing McKO and NcKO to their appropriate littermate EP2+ controls removed genetic variation, a strong modifier of seizure susceptibility and SE-induced phenotypes (14, 15).

Beneficial behavioral outcomes have been attributed to global EP2 ablation in a mouse model of amyotrophic lateral sclerosis (16). Myeloid cell ablation of EP2 suppressed inflammation, enhanced amyloid clearance, and prevented synaptic injury and memory deficits in an Alzheimer’s disease model (17, 18). Moreover, multiple beneficial effects are encountered after administration of a brain-permeable EP2 antagonist starting 2-4 hours after SE onset, namely a reduction in delayed mortality, blunted neuroinflammation, prevention of BBB opening, neuroprotection, accelerated weight regain, and cognitive improvement (3, 8-10). In contrast, global EP2 ablation increased infarct size in a model of focal ischemia (19), and we have shown that intraventricular delivery of an EP2 agonist immediately after pilocarpine-induced SE in a rat model is neuroprotective (7). These seemingly contrasting observations might be attributed to divergent consequences of EP2 activation after neurological insult, wherein EP2 activation is neuroprotective early and neurotoxic later.

Disruption of the BBB is a common event in numerous neurological disorders and contributes to brain injury (20). Leakage of serum albumin into the brain after SE could promote the development of epilepsy because direct administration of albumin into the brain can result in the appearance of an epileptic focus that evolves into spontaneous seizures (13). Additionally, deterioration of the BBB has been documented in humans after seizures by measuring the extravasation of serum albumin into the brain parenchyma (2). In rodents the integrity of the BBB is compromised within hours of SE and persists for weeks (21, 22).

A surprising finding in the current studies was the elevated cortical albumin levels in saline treated McKO mice. The elevated baseline levels of cortical albumin in the McKO mice might be attributed to a protective effect of myeloid cell EP2 under basal conditions. The finding that the BBB is leaky in naïve saline treated McKO mice is in seeming disagreement with our previous findings showing that EP2 antagonism for two days after SE did not facilitate albumin entry into the cortex (8). One explanation might be that silencing EP2 signaling in myeloid cells for up to 10 weeks during development as occurs in the McKO mice, as opposed to two days via EP2 antagonism, contributes to BBB deterioration.

Increased permeability of the BBB is closely correlated with a neuroinflammatory response typified by expression of inflammatory mediators in brain and cerebral vessels (23). SE results in the rapid induction of numerous inflammatory cytokines and chemokines with IL-1β and Cox-2 induction occurring within 30 minutes of SE onset, followed by induction of Ccl2, TNFα, and IL-6 (3). Pharmacologic EP2 antagonism after SE results in broad quenching of inflammatory mediators and maintenance of BBB integrity (8). Disrupting EP2 signaling in myeloid cells selectively attenuated the induction of the pro-inflammatory cytokine IL-6 in hippocampus four days after pilocarpine-induced SE. This finding suggests that IL-6 production is induced in an EP2-dependent manner, whereas more complex signaling pathways might be responsible for the other mediators and glial markers. Interestingly, IL-6 has been implicated in BBB dysfunction in a fetal ovine brain ischemia model (24). Thus, protection of the BBB integrity in McKO mice after SE might be related to dampened IL-6 induction. IL-6 also promotes epileptogenesis induced by murine viral encephalitis (25), and IL-6 induces epileptiform discharges in vitro and spontaneous seizures in C57BL/6 mice (26). These findings raise the question whether either EP2 antagonism or conditional ablation of EP2 from myeloid cells prevents the development of spontaneous seizures, which is underway in our laboratory.

Elevated expression of TNFα and Ccl2 was detected in hippocampus of NcKO mice after SE, compared to nEP2+ mice. TNFα can be both pro- and anti-seizure through its receptors TNFR1 and TNFR2, respectively (27, 28). The chemokine Ccl2 is a ligand for the chemokine receptor Ccr2, and Ccl2 signaling is involved in peripheral monocyte recruitment to the brain (29) and regulation of BBB permeability (30). Peripheral monocyte recruitment to the brain is pathogenic after SE, enhancing neuronal damage, limiting weight regain, and eroding the BBB (11). Thus, the deleterious effects of SE in NcKO mice could be attributed to increased monocyte recruitment to the brain. However, additional work is required to test this idea.

Microglial EP2 has been implicated in paracrine neurotoxicity following lipopolysaccharide (LPS)-induced inflammation *in vitro* (31). Our own studies have demonstrated that systemic EP2 antagonism, delayed for 2-4 hr after SE, is neuroprotective (8, 9). In the current studies, ablation of EP2 from myeloid cells did not result in neuronal protection after SE, as neuronal damage was nearly equal in McKO and littermate mEP2+ mice. These conflicting findings might be attributed to the different inflammatory stimuli, LPS vs. SE, acting on myeloid immune cells. Even more surprising was the enhancement of neuronal damage in NcKO mice compared to control nEP2-sufficient mice. Thus, it currently remains unclear on which cell type EP2 antagonism elicits neuroprotective effects.

In man, mortality is high during refractory SE that requires general anesthesia (32, 33), and the 30-day mortality is ∼35% in adults who experience at least an hour of SE (34). Mortality is also encountered in rodents experiencing SE, and mortality rate is positively correlated with time spent in SE (35). We found no role for myeloid cell EP2 or neuronal EP2 in acute mortality during SE. In contrast, systemic EP2 antagonism in mice decreased mortality in the seven days following SE (3, 8). In patients, delayed mortality after SE is associated with gradual or abrupt cardiac decompensation (36). Thus, it is possible that reduced delayed mortality observed after systemic EP2 administration might be attributed to antagonism of non-myeloid or peripheral (e.g., cardiac) EP2 receptors. However, conditional ablation of COX-2 limited to forebrain neurons also rescued delayed mortality (7), suggesting the relevant EP2 receptors are central rather than peripheral. Additional work is needed to identify the EP2 receptors responsible for enhanced mortality after SE.

As an additional assessment of behavioral deficits after SE we performed a modified Irwin test, in which we judged pilocarpine-treated mice over a number of parameters in the four days following SE. Notably, McKO mice performed significantly better than mEP2+ mice with relieved ptosis playing a dominant role in the rescued neurobehavioral deficits, whereas NcKO mice showed worsened ptosis, poor body posture, and altered activity. Ptosis in human patients can be associated with cerebral strokes, with bilateral ptosis typically seen in those with large cerebral infarctions (37). In the current study, ptosis was correlated with enhanced BBB erosion in McKO mice entering SE. However, IgG extravasation into the cortex and brain Prussian blue (hemosiderin) staining was not noticeable (not shown), suggesting some selectively to pilocarpine-induced BBB permeability.

It is currently unclear why neuronal and myeloid EP2 signaling results in opposing outcomes after SE. EP2 activates downstream signaling either by a G protein-coupled or a G protein-independent mechanism. G protein coupled signaling through Gαs elevates cAMP levels, which can either activate protein kinase A (PKA) (38) or exchange protein active (Epac) (39, 40). Alternatively, EP2 activation can signal through β-arrestin in a G protein-independent pathway. Each EP2-dependent signaling cascade can result in divergent cellular consequences (41, 42). Thus, the current results might be attributed to one EP2-driven signaling pathway dominating over the other signaling pathways in a cell type-specific manner, highlighting the complexities of neuroinflammatory signaling in the brain after insults. Nevertheless, our findings reinforce the notion that prostaglandin receptor EP2 should continue to be explored as a therapeutic target to treat SE, but also caution us to target EP2 in a cell-specific manner.

## Materials and Methods

### Mice

All mice used in the current study were housed in the same rooms in Emory’s animal facility during the course of the experiment, with one room designated for animal breeding and maintenance and another room solely for SE experiments. Lighting was on a 12-hour on/off cycle, and mice are fed and watered *ab libitum*. We obtained male *Cd11b-Cre* (43) and female *EP2*^*flox/flox*^ (18) from K. Andreasson (Stanford University, CA) maintained on the C57BL/6 (Charles River) background. The mice were subsequently intercrossed to propagate the colony. *Cd11b-Cre;EP2*^*flox/flox*^ mice were identified as EP2 conditional knockout mice (McKO), whereas EP2-sufficient littermates (mEP2+) were either homozygous *EP2*^*flox/flox*^ or heterozygous *EP2*^*+/flox*^ and negative for the *Cd11b-Cre* transgene. Of all 170 mice generated, 13.5% of the offspring were male and McKO, and 28.2% were male and mEP2+, in line with the expected Mendelian ratios. We have selected the CD11b-Cre driver to ablate EP2 from myeloid cells based on the following considerations. First, the myelomonocytic marker CD11b is expressed by both microglia and monocytes. Second, we and others have demonstrated the involvement of both microglia and monocytes in the ensuing neuroinflammatory response following SE (8, 11, 44).

In separate matings, female mice expressing Cre recombinase under control of the synapsin 1 promoter (7, 45) were bred to male *EP2*^*flox/flox*^ mice (18) to generate females expressing Cre and heterozygous for floxed EP2. These females were further bred with heterozygous floxed EP2 males to generate EP2-sufficient littermates (nEP2+), either homozygous *EP2*^*flox/flox*^ or heterozygous *EP2*^*+/flox*^ and negative for the s*ynapsin 1-Cre* transgene and neuron-specific conditional EP2 conditional knockouts (NcKO). Of all 210 mice generated, 11.4% of the offspring were male and NcKO, and 24.8% were male and mEP2+. We have previously utilized synapsin-1-Cre to robustly ablate Cox-2 from principal forebrain neurons (7).

Male mice were utilized in the current studies as, after pilocarpine, male mice more readily entered SE when compared to female littermates; often the pilocarpine dose required to elicit SE was lethal in female mice. Attempts were made to validate three commercial EP2 antibodies using tissues from EP2 knockout mice (46) obtained from The Jackson Laboratories, or tissues from another EP2 knockout line (47) obtained from Dr. Tomohiro Aoki (Department of Molecular Pharmacology, Research Institute, National Cerebral and Cardiovascular Center, Suita, Osaka, Japan).

### Pilocarpine Injection

To minimize peripheral side effects of pilocarpine, mice were injected intraperitoneally (i.p.) with methylscopolamine and terbutaline (2 mg/kg each in 0.9% saline solution). After 20 minutes, pilocarpine hydrochloride (328 mg/kg in 0.9% saline, freshly prepared) was injected i.p. to induce SE. Control mice of each genotype received methylscopolamine and terbutaline, followed by injection of saline solution instead of pilocarpine. Seizures were classified as previously described (11, 15) and scored every 5 min as follows: a score of 0 represents normal behavior (walking, exploring, sniffing, grooming); a score of 1 represents immobility, staring, jumpy/cured-up posture, a score of 2 represents automatisms (repetitive blinking, chewing, head bobbing, vibrissae twitching, scratching, face-washing, “star-gazing”); a score of 3 represents partial body clonus, occasional myoclonic jerks, shivering; a score of 4 represents whole-body clonus, “corkscrew” turning and flipping, loss of posture, rearing and falling; a score of 5 (SE onset) represents nonintermittent seizure activity consisting of stage 3 or 4 repeatedly; a score of 6 represents wild running, bouncing, and tonic seizures; and a score of 7 represents death. After one hour, SE was interrupted by diazepam (10 mg/kg, i.p.). Mice were fed moistened rodent chow, monitored daily, and injected with 5% (wt/vol) dextrose in lactated Ringer’s solution (0.5 mL i.p.) when necessary. In tests of McKO and NcKO mice, their respective age-matched littermates were treated with pilocarpine or saline alongside the conditional knockout mice, and tissue was processed together.

### Modified Irwin test

A modified Irwin test (48) was performed to assess the health and well-being of mice after pilocarpine-induced SE. The test comprised seven parameters (ptosis, lacrimation, body posture, tremors, body dragging, hyper- or hypoactivity, coat appearance) that can be measured simply by experimenter observation. The test was given three times: 48, 72, and 96 hours after SE onset. Each parameter was scored on a three-point scale (i.e., 0 = normal, 1 = mild to moderate impairment, and 2 = severe impairment) with a total score ranging from 0 to 14. A total score of 0 as a sum of all 7 parameters indicates a normal healthy mouse. A total score ranging from 1 to 14 indicates a mouse that is impaired.

### Tissue Processing

Four days after SE onset, mice were anesthetized deeply with isoflurane, perfused through the heart with PBS solution, and their brains rapidly removed from the cranium. The brain was immediately bisected through the midline, and the left hemisphere was fixed for 24 hours in 4% (wt/vol) paraformaldehyde at 4°C for Fluoro-Jade B staining. The hippocampus and cortex were dissected from the right half of the brain, immediately frozen on dry ice and stored at −80°C for RNA isolation and Western blot analysis, respectively.

### Western Blot Analysis of Albumin

Cerebral cortices from saline perfused mice were homogenized in 10 volumes of RIPA buffer with protease and phosphatase inhibitors (Thermo Scientific). Brain homogenates were subsequently sonicated to shear DNA then centrifuged at 14,000 x *g* for 30 minutes at 4°C to remove nuclei and cell debris. Brain protein was run on a 4-20% Mini-PROTEAN TGX Gel and electroblotted onto PVDF membranes (Millipore).

Membranes were blocked for one hour at room temperature with Odyssey blocking solution (Li-Cor), then incubated overnight at 4°C with primary antibodies for albumin (1:1000; Cell Signaling) and GAPDH (1:10,000; Calbiochem), followed by incubation with polyclonal IRDye secondary antibodies 680LT and 800CW (1:15,000; Li-Cor). The blots were imaged by using Li-Cor imaging systems using channels 700 and 800. Because all samples could not be run on the same gel, one sample was run on each blot for interblot normalization. Band intensity was measured and corrected for nearby background intensity by the Li-Cor software. The log value of the albumin/GAPDH ratio for each sample was then determined. The average log ratio for selected groups was compared by ANOVA with post hoc Sidak adjustment. The fold-change for each group was plotted on a logarithmic scale.

### Invalidation of EP2 Antibodies

In an effort to monitor EP2 expression in the myeloid EP2 knockout mouse, we initially turned to commonly available antibodies. To validate these antibodies we compared western blots from tissues of wild-type and EP2 knockout mice. Brain, heart, uterus, and kidney from perfused global EP2 KO mouse organs were kindly provided by Dr. Tomohiro Aoki (Department of Molecular Pharmacology, Research Institute, National Cerebral and Cardiovascular Center, Suita, Osaka, Japan) (47). The tissues were homogenized and centrifuged as described above. Membranes were blocked for one hour at room temperature with Odyssey blocking solution (Li-Cor), then incubated overnight at 4°C with primary rabbit polyclonal antibodies that are directed to the remaining EP2 coding sequence from *EP2* knockout mice (LS-C406698, 1:200 or LS-B6048, 1:750 or LS-C200473, 1:750, all from LifeSpan Biosciences, Inc.) and GAPDH (1:10,000; Calbiochem), followed by incubation with polyclonal IRDye secondary antibodies 680LT and 800CW (1:15,000; Li-Cor). The blots were imaged by using Li-Cor imaging systems using channels 700 and 800. Both polyclonal antibodies LS-C200473 (Figure S1A) and LS-B6048 (Figure S1B) detected a protein migrating at approximately 42 kDa in brain, uterine, and kidney homogenates obtained from EP2 knockout mice. These two antibodies detected no discernible proteins of lower molecular weight. Finally we found that polyclonal antibody LS-C406698 detected a protein migrating at ∼60 kDa in kidney and uterine homogenates, but not in homogenates of heart and brain, obtained from EP2 sufficient mouse tissue (Figure S1C). In addition a band of similar size was detected in BV-2 cells stably expressing human EP2 as well as parent BV-2 cells that lack functional endogenous mouse EP2. In addition we also observed a band migrating near the 37 kDa marker in all tissue and cellular homogenates, including the EP2 knockout brain. We conclude these three polyclonal antibodies are unreliable.

### Fluoro-Jade B Labeling

After 24 hours in 4% paraformaldehyde the left brain hemisphere was cryoprotected in 30% (wt/vol) sucrose in PBS solution. Brains were subsequently frozen in 2-methylbutane and sectioned coronally at 25 μm by using a freezing/sliding microtome. Every 12^th^ section for a total of approximately five to six sections from each mouse hemisphere was used for Fluoro-Jade B staining to label degenerating neurons. Sections were mounted on Superfrost Plus Microscope Slides (Fisher Scientific) and allowed to air-dry overnight. Sections were immersed in 0.06% potassium permanganate for 15 minutes with gentle agitation, rinsed for one minute in distilled water, and then transferred to the Fluoro-Jade B staining solution (0.0001% wt/vol FJB in distilled water with 0.1% acetic acid) for 30 minutes with gentle agitation. Sections were rinsed with three one-minute changes of distilled water and air-dried. The slides were immersed in xylene and then coverslipped with D.P.X. mounting media (Sigma-Aldrich). Sections between bregma −1.34 and −2.40 were examined with a fluorescent microscope.

Following Fluoro-Jade B staining, images were obtained from three hippocampal areas (hilus, CA1, CA3) in each of the sections. Two researchers unaware of mouse genotype and experimental conditions counted the number of Fluoro-Jade B-positive neurons in the hippocampus. Only positive neurons with a near-complete cell body shape and size were tabulated. Cell counts were expressed as the total number of Fluoro-Jade B-positive cells per section for each region. The inter-rater reliability was determined by measuring pairwise correlation of the average number of Fluoro-Jade B-positive neurons countered in each animal in the study. The Pearson coefficient was 0.99 with a slope of 0.92.

### qRT-PCR

Total RNA from mouse hippocampus was isolated by using TRIzol (Invitrogen) with the PureLink RNA Mini Kit (Invitrogen). RNA concentration and purity were measured by A260 value and the A260/A280 ratio, respectively. We typically recovered 5-15 μg RNA from each hippocampus with A260/280 ratio = 2.1-2.2. First-strand cDNA synthesis was performed with 1.0 μg of total RNA, using qScript cDNA SuperMix (#95048, Quantabio) in a reaction volume of 20 μL at 25°C for five minutes, then 42°C for 30 minutes. The reaction was terminated by heating at 85°C for five minutes. qRT-PCR was performed by using eight μL of 50x-diluted cDNA, 0.1-0.5 μM of primers, and 2x iQ SYBR Green Supermix (Bio-Rad Laboratories), with a final volume of 20 μL, in the iQ5 Multicolor Real-Time PCR Detection System (Bio-Rad). Cycling conditions were as follows: 95°C for two minutes followed by 40 cycles of 95°C for 15 seconds and then 60°C for one minute. Melting curve analysis was used to verify single-species PCR product. Fluorescent data were acquired at the 60°C step. The geometric mean of the cycle thresholds for β-actin, GAPDH, and HPRT1 was used as internal control of relative quantification (Tables S1 and S2). Samples without cDNA samples served as the negative controls. We also examined efficiency of the PCR reaction by determining the Ct values for serial dilutions of our cDNA and plotting the resulting Ct values as a function of the negative (log dilution factor) (Figure S2). A PCR reaction that is 100% efficient would yield a resulting slope of 3.33; our mean efficiency was 94.6±1.7% with a range of 83-105%.

### Splenocyte Isolation and Flow Cytometry

Spleens were harvested from mEP2+ and McKO mice. Splenocytes were obtained by passing the spleen contents twice through a 70 μm mesh with the aid of a syringe handle followed by lysing the red blood cells with AKC lysing buffer (Thermo Fisher Scientific). Splenocytes were then washed with cold PBS containing 5% FBS. Single-cell suspensions were stained with anti-CD11b-PerCP (M1/70; BioLegend) antibodies. Cells were sorted on a FACSAria II (BD) running Diva6. Data were analyzed with FlowJo software (Tree Star).

### RNA Extraction and Amplification from Sorted CD11b+ Splenocytes

RNA was extracted from isolated splenocytes using Quick-RNA MiniPrep (Zymo) according to manufacturer’s instructions. RNA samples were analyzed with the BioAnalyzer (Agilent Technologies, Inc.). Only samples with an RNA integrity value of at least seven were selected for further analysis. RNA from all six mice met this criterion with a mean value of 8.95. One round of poly-A RNA amplification was performed on each sample by using Message *BOOSTER* cDNA synthesis kit for quantitative PCR (Lucigen) following manufacturer’s instructors. Gene-expression analysis was performed as described above. The cycle threshold value for CD11b was used as internal control of relative expression in myeloid cells.

### Data and Statistical Analysis

The results are expressed as mean values ± SEM. Statistical analysis was performed using GraphPad version 6 (GraphPad Software). One- or two-way ANOVA was used for comparison of selected means with Sidak test where appropriate. Unpaired t-tests were used for comparing two groups. For analysis of gene induction, the mean ΔΔCt values were compared between selected groups. After statistical analysis, individual ΔΔCt values from each sample group were converted to fold-change by 2^ΔΔCt, and the fold-change from each group was plotted on a logarithmic scale. For all analyses, the differences were considered to be statistically different if p<0.05.

## Acknowledgements

We thank K. Andreasson (Stanford University, CA) for sharing EP2 flox mice and CD11b-Cre mice, T. Aoki (National Cerebral and Cardiovascular Center, Suita, Osaka, Japan) for EP2 knockout tissues, and A. Rea (Emory University, GA) for assistance with flow sorting. This research project was supported in part by Emory University Pediatric Flow Cytometry Core and Emory University Genomics Core as well as National Institutes of Health, Office of the Director, National Institute of Neurological Disorders and Stroke Grant R01 NS097776 (to R.D.).

## Supplementary Figures and Tables

**Figure S1.**
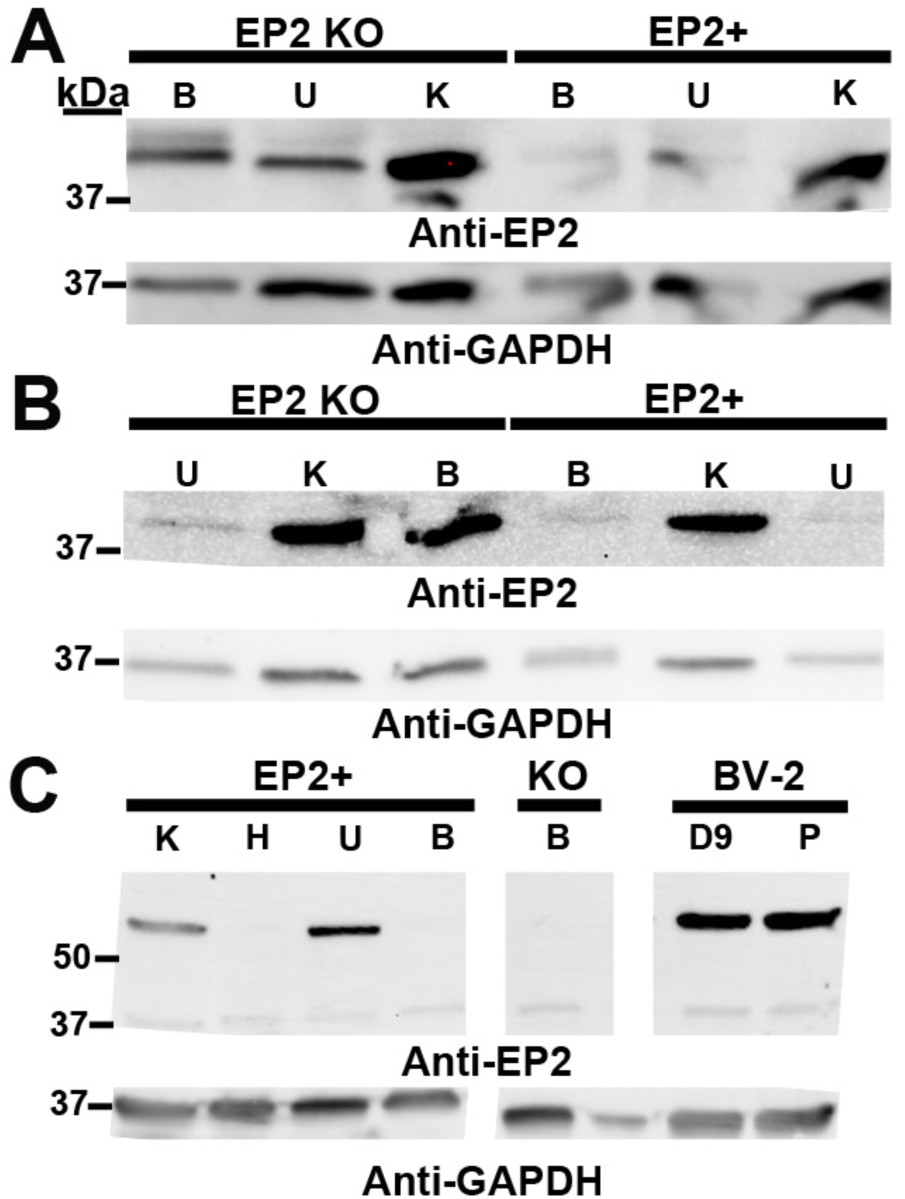
Unsuccessful attempt to validate three EP2 antibodies. (A) Polyclonal EP2 antibody (LS-C200473 from LS-Bio) was tested in samples obtained from EP2+ and global EP2 knockout (KO) animals from Japan. Approximately 77% of the EP2 coding sequence had been replaced by a neo cassette (through T281), and the antibody was chosen to minimally overlap the potential remaining EP2 protein sequence. A band of approximately 42 kDa was observed in brain (B), uterine (U), and kidney (K) homogenates obtained from EP2+ mice. A corresponding band was also observed in brain, uterine, and kidney homogenates from KO mice. (B) A band of approximately 42 kDa was observed in uterine, kidney, and brain homogenates obtained from EP2 KO mice from Japan by another polyclonal EP2 antibody (LS-B6048). A corresponding band was also detected in kidney homogenates from EP2+ mice, whereas a lighter band at approximately the same size was observed in brain and uterine homogenates. (C) Polyclonal EP2 antibody (LS-C406698) detected a protein of ∼ 60 kDa in kidney and uterine homogenates, but not in heart (H) and brain homogenates from EP2+ mice. No band of similar size was detected in EP2 KO mice (KO) from Jackson laboratories. However, a similar molecular weight band was detected in BV-2 cells stably expressing human EP2 (D9), and a similar intensity band was observed in parent EP2 BV-2 cells that lack functional EP2 (P). All three panels were obtained from the same blot and imaged under identical conditions. Superfluous lanes were removed for data presentation. GAPDH was used in each blot as loading control.

**Figure S2.**
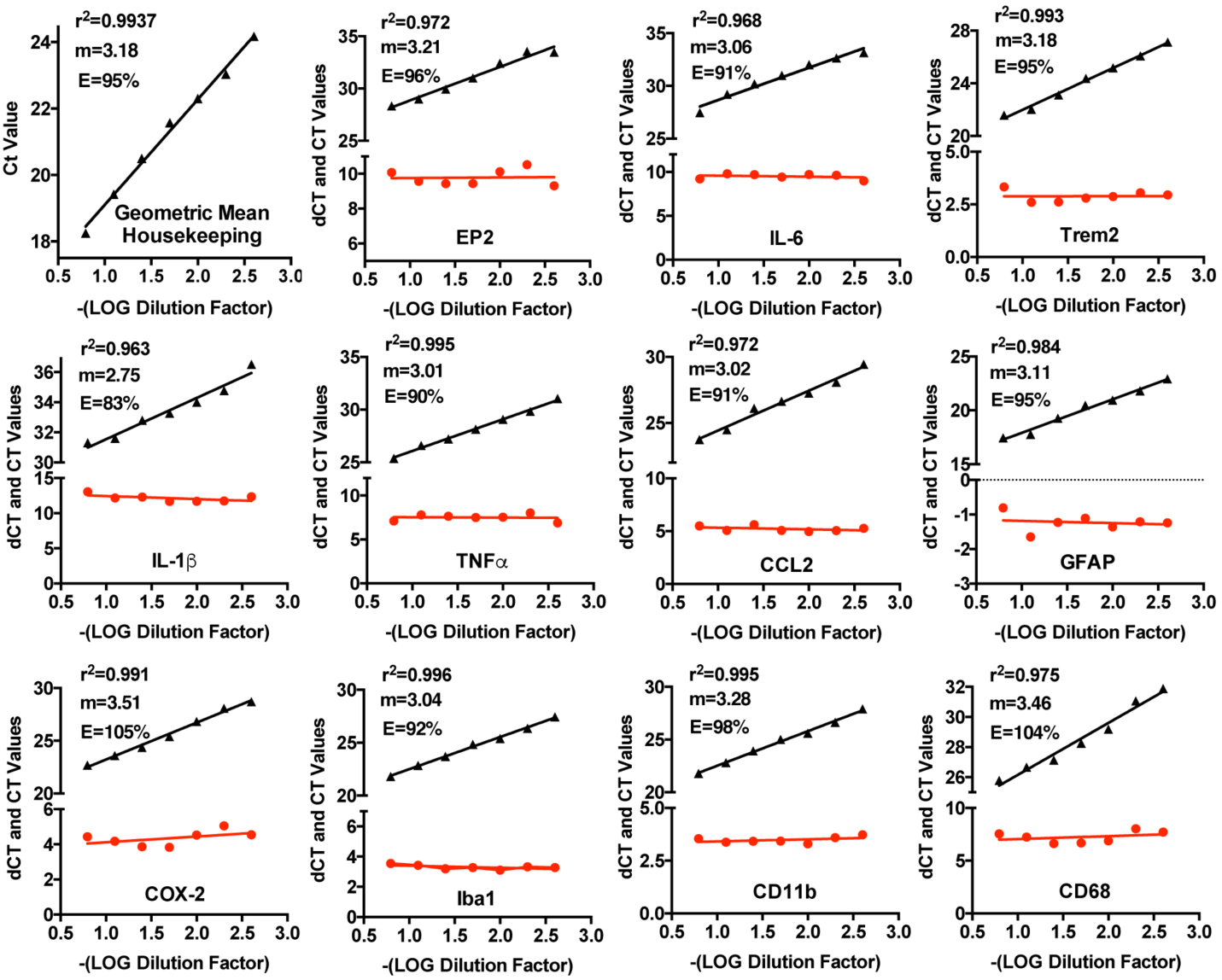
Standard curves for verification of qRT-PCR. Quantitative PCR was performed on serial dilutions of hippocampal cDNA prepared from a mEP2+ mouse, and the Ct value for each transcript was plotted for each dilution (black triangles with black lines). Each primer set used in the study was assayed as well as the three house keeping genes (GAPDH, β-actin, and HPRT1). The geometric mean of the three house keeping genes is plotted at the upper left. The delta Ct values for each gene were also determined for each gene target by subtracting the geometric mean of the housekeeping Ct value from the target Ct value obtained from the corresponding dilution. The delta Ct values for each target were also plotted (red circles and red lines).

**Table S1.** CT values and geometric means of housekeeping genes in four groups (n=8 mEP2+ basal, n=8 McKO basal, n=8 mEP2+ pilocarpine, and n=8 McKO pilocarpine) from mice four days after treatment with saline or pilocarpine.

| Genes | mEP2+<br>basal | McKO<br>basal | mEP2+<br>pilocarpine | McKO<br>pilocarpine |
| --- | --- | --- | --- | --- |
| HPRT1 | 23.6±0.2 | 23.8±0.4 | 24.0±0.3 | 23.8±0.2 |
| β-actin | 22.0±0.3 | 21.6±0.2 | 21.4±0.3 | 20.8±0.3 |
| GAPDH | 18.1±0.1 | 17.9±0.1 | 18.3±0.2 | 17.9±0.1 |
| **Geomean** | **21.1±0.2** | **20.9±0.2** | **21.1±0.2** | **20.7±0.2** |

Data are the mean ± SEM.

**Table S2.** CT values and geometric means of housekeeping genes in four groups (n=8 nEP2+ basal, n=8 NcKO basal, n=8 nEP2+ pilocarpine, and n=8 NcKO pilocarpine) from mice four days after treatment with saline or pilocarpine.

| Genes | nEP2+<br>basal | NcKO<br>basal | nEP2+<br>pilocarpine | NcKO<br>pilocarpine |
| --- | --- | --- | --- | --- |
| HPRT1 | 26.4±0.5 | 26.8±0.5 | 27.4±0.5 | 27.6±0.6 |
| β-actin | 23.3±0.6 | 23.5±0.8 | 23.4±0.3 | 22.3±0.7 |
| GAPDH | 19.0±0.3 | 19.5±0.8 | 20.0±0.5 | 18.4±0.4 |
| **Geomean** | **22.7±0.4** | **23.0±0.6** | **23.4±0.3** | **22.4±0.5** |

Data are the mean ± SEM.

